# ALIBY: ALFA Nanobody-Based Toolkit for Imaging and Biochemistry in Yeast

**DOI:** 10.1101/2022.07.18.500560

**Authors:** Dipayan Akhuli, Anubhav Dhar, Aileen Sara Viji, Bindu Bhojappa, Saravanan Palani

## Abstract

Specialized epitope tags continue to be an integral component in various biochemical and cell biological applications such as fluorescence microscopy, immunoblotting, immunoprecipitation, and protein purification. However, until recently, no single tag could offer this complete set of functionalities on its own. Here, we present a plasmid-based toolkit named ALIBY (**AL**FA Toolkit for **I**maging and **B**iochemistry in **Y**east) that provides a universal workflow to adopt the versatile ALFA tag/^Nb^ALFA system within the well-established model organism *Saccharomyces cerevisiae.* The kit comprises of tagging plasmids for labelling a protein-of-interest with the ALFA tag, and detection plasmids encoding a fluorescent protein-tagged ^Nb^ALFA for live-cell imaging purposes. We demonstrate the suitability of ALIBY for visualizing the spatiotemporal localization of yeast proteins (i.e., cytoskeleton, nucleus, centrosome, divisome and exocyst) in live cells. Our approach has yielded an excellent signal-to-noise ratio without off-targeting or effect on cell growth. In summary, our yeast-specific toolkit aims to simplify and further advance the live-cell imaging of differentially abundant yeast proteins while also being suitable for biochemical applications.

**Importance:** In yeast research, conventional fluorescent protein tags and small epitope tags are widely used to study the spatiotemporal dynamics and activity of proteins. Though proven to be efficient, these tags lack the versatility for usage across different cell biological and biochemical studies of a given protein-of-interest. Therefore, there is an urgent need for a unified platform for visualization, biochemical, and functional analyses of proteins-of-interest in yeast. Herein, we have engineered ALIBY, a plasmid-based toolkit which expands the benefits of the recently developed ALFA tag/^Nb^ALFA system to studies in the well-established model organism *Saccharomyces cerevisiae.* We demonstrate that ALIBY provides a simple and versatile strain construction workflow for long duration live-cell imaging and biochemical applications in yeast.

## Introduction

*Saccharomyces cerevisiae* is a well-established model system for studying basic principles of eukaryotic cell biology. It is a simple, unicellular eukaryote that permits easy genetic modifications such as gene deletions or tagging using a standard PCR-based strategy(1). Numerous epitope tags have been designed to aid in investigating protein localization and performing biochemical analysis. MBP and GST tags have been used for affinity purification of fusion proteins but can impose heavy metabolic load when overexpressed(2). FLAG-, HA- and myc-tags are utilized in immunostaining, immunoprecipitation (IP), Western blotting, and protein purification experiments but their large antibody partners make them unsuitable for super-resolution microscopy(3). The HA-tag is reported to lose its immunoreactivity during apoptosis, limiting its usability in cell death-related studies(4). SPOT and EPEA tags perform well in super-resolution microscopy but their nanobody partners show non-specific binding and have poor binding affinity(5–7). Additionally, live-cell imaging has not been demonstrated for these tags(3). A cause of concern is variability in dynamics of any given protein tagged with different epitope tags for different purposes. The field of yeast cell biology currently lacks a universal tag that can cover a wide range of biological applications.

The recently developed ALFA tag excels at all of these functionalities simultaneously(3), and has been successfully employed in bacterial and mammalian systems(8–12). Here, we aim to extend the ALFA/^Nb^ALFA system to the model organism *Saccharomyces cerevisiae* via a plasmid-based toolkit named ALIBY. The kit consists of two sets of plasmids: 1) Tagging plasmids for labelling a protein-of-interest (POI) at its native genomic locus with a C-terminal ALFA tag, and 2) Detection plasmids expressing ^Nb^ALFA tagged to a fluorescent protein under various promoters for *in vivo* detection and visualization of the ALFA-tagged POI. ALIBY can streamline the process of yeast strain construction for diverse biochemistry and cell biology studies, making it less laborious and expensive. To this end, we demonstrate the functionality of ALIBY for live-cell imaging, Western blotting, and immunoprecipitation of various cytoskeletal and other proteins in *Saccharomyces cerevisiae*.

## Results

### Design of plasmids

We engineered two families of plasmids as a part of ALIBY: 1) Tagging plasmids, and 2) Detection plasmids. The tagging plasmids are based on the pYM series of plasmids(13), and serve as a template for amplification of the ALFA tag-containing PCR cassette using the universal S2/S3 primer set(1, 13) (**Fig. 1a, Table 1**). This PCR cassette, containing one of four selectable markers of choice *(kanMX4, hphNT1, natNT2* and *HIS3MX6),* can be used to fuse the ALFA tag to the C-terminal of any POI via a commonly used homologous recombination-based strategy(1).The detection plasmids are based on the pRS30-series of yeast integrative plasmids (YIp)(14), and they express ^Nb^ALFA linked to mNeonGreen via a flexible 40 amino acid linker (40aaL) at its C-terminus (**Fig. 1b, Table 1**). mNeonGreen was selected because it is brighter, has a higher quantum yield, is more photostable and has a lower maturation time than commonly used green and yellow fluorescent proteins(15).The 40 amino acid linker has been used as a spacer between proteins and their fused fluorescent tags such that they do not interfere with the folding and normal functioning of the POI(16).The ^Nb^ALFA-Linker-mNeonGreen (hereafter referred to as ^Nb^ALFA-L-mNG) construct has been designed with the choice of 4 different constitutive promoters – *PCYC1, PADH1, PTEF,* and *PGPD* (in increasing order of promoter strength(17)) in combination with 4 different vector backbones of the pRS30-series containing *HIS3, TRP1, LEU2* and *URA3* selectable markers (**Table 1**). Thus, to construct a strain with a POI C-terminally fused to the ALFA tag, a PCR cassette is generated using a tagging plasmid as template with genespecific S2/S3 primers, which is then transformed into a starting strain of choice. This strain can be used for biochemical purposes like immunoblotting, IP, CoIP, protein purification, mass spectrometry, and also fixed-cell imaging using immunofluorescence and superresolution microscopy. Additionally, a linearized detection plasmid containing a marker of choice and expressing the ^Nb^ALFA-L-mNG construct under a suitable promoter depending on the experimental design, can be integrated into the above strain carrying the ALFA tag. This strain can now be used for live-cell imaging purposes (**Fig. 1c**). The aforementioned workflow significantly simplifies the strain construction process by saving time and resources and making it less laborious and extensive.

**Fig. 1.**
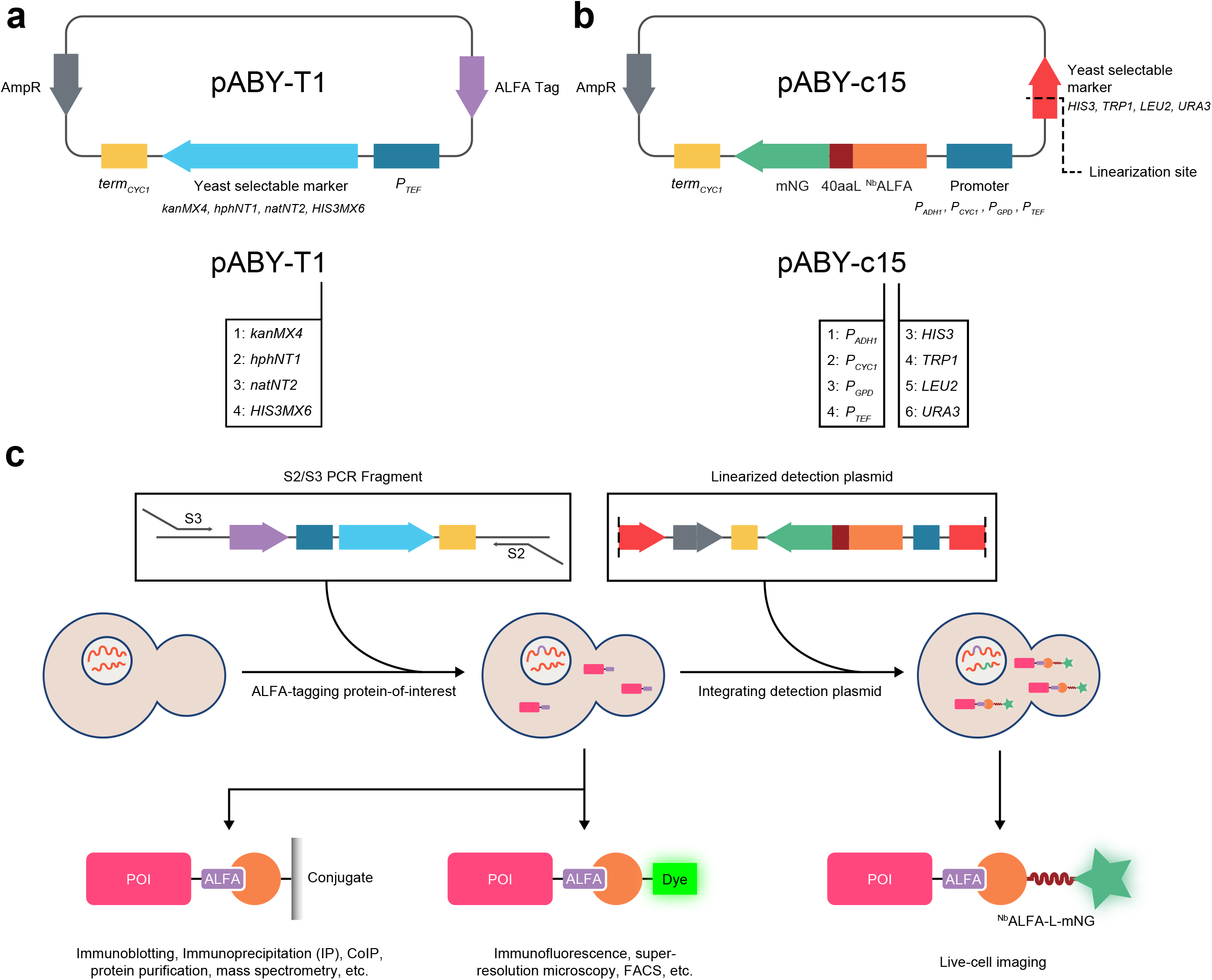
Design of detection and tagging plasmids comprising the toolkit and the proposed workflow. **a** (Top) Schematic map of a representative tagging plasmid and (bottom) its naming scheme. The number in the plasmid name code corresponds to the yeast selectable marker as tabulated. **b** (Top) Schematic map of a representative pRS-based detection plasmid and (bottom) its naming scheme. 40aaL – 40 amino acid linker; mNG – mNeonGreen. AmpR encodes a beta-lactamase responsible for conferring resistance to ampicillin, used as bacterial selectable marker. The black dashed line marks the linearization site in the yeast selectable marker gene. In the plasmid name code, the first digit corresponds to the promoter while the second digit corresponds to the yeast selectable marker, as tabulated. **c** Schematic of the recommended workflow for strain construction. S2 primer overhang sequence: 5’-ATCGATGAATTCGAGCTCG-3’; S3 primer overhang sequence: 5’-CGTACGCTGCAGGTCGAC-3’. Starting from a particular strain of choice (middle left), the PCR fragment generated from the tagging plasmid using S2/S3 primers (top left) can be transformed to generate the tagged strain (middle centre). The linearized detection plasmid (top right) containing a promoter and a selectable marker of choice can now be integrated into the tagged strain to obtain the nanobody-containing tagged strain (middle right), which can be used for live-cell imaging (bottom right). The colored blocks in the top row correspond to those already labelled in panels **a** and **b**. Color code for schematics in bottom row: POI (pink), ALFA tag (purple), ^Nb^ALFA (orange), dye (fluorescent green), 40 amino acid linker (brown), mNeonGreen (green).

**Table 1:** Details of tagging and detection plasmids.

| Plasmid No. | Description | Name Code | Backbone | Promoter | Selectable marker | Linearization site | PCR Fragment size (bp) |
| --- | --- | --- | --- | --- | --- | --- | --- |
| piSP532 | pYM14-ALFA-*kanMX4* | pABY-T1 | pFA6a | - | *kanMX4* | - | 1521 |
| piSP534 | pYM16-ALFA-*hphNT1* | pABY-T2 | pFA6a | - | *hphNT1* | - | 1750 |
| piSP536 | pYM17-ALFA-*natNT2* | pABY-T3 | pFA6a | - | *natNT2* | - | 1367 |
| piSP538 | pFA6a-ALFA-*HIS3MX6* | pABY-T4 | pFA6a | - | *HIS3MX6* | - | 1365 |
| piSP551 | pRS305-*P<sub>ADH1</sub>*<sup>Nb</sup>ALFA-L-mNG-*term<sub>CYC1</sub>* | pABY-c15 | pRS305 | *P<sub>ADH1</sub>* | *LEU2* | KasI | - |
| piSP553 | pRS305-*P<sub>CYC1</sub>*<sup>Nb</sup>ALFA-L-mNG-*term<sub>CYC1</sub>* | pABY-c25 | pRS305 | *P<sub>CYC1</sub>* | *LEU2* | KasI | - |
| piSP555 | pRS305-*P<sub>GPD</sub>*<sup>Nb</sup>ALFA-L-mNG-*term<sub>CYC1</sub>* | pABY-c35 | pRS305 | *P<sub>GPD</sub>* | *LEU2* | KasI | - |
| piSP557 | pRS305-*P<sub>TEF</sub>*<sup>Nb</sup>ALFA-L-mNG-*term<sub>CYC1</sub>* | pABY-c45 | pRS305 | *P<sub>TEF</sub>* | *LEU2* | KasI | - |
| piSP765 | pRS303-*P<sub>TEF</sub>*<sup>Nb</sup>ALFA-L-mNG-*term<sub>CYC1</sub>* | pABY-c43 | pRS303 | *P<sub>TEF</sub>* | *HIS3* | PstI | - |
| piSP767 | pRS304-*P<sub>TEF</sub>*<sup>Nb</sup>ALFA-L-mNG-*term<sub>CYC1</sub>* | pABY-c44 | pRS304 | *P<sub>TEF</sub>* | *TRP1* | SnaBI | - |
| piSP565 | pRS306-*P<sub>TEF</sub>*<sup>Nb</sup>ALFA-L-mNG-*term<sub>CYC1</sub>* | pABY-c46 | pRS306 | *P<sub>TEF</sub>* | *URA3* | NdeI | - |

### Characterization of plasmids

To characterize the two sets of plasmids, we first constructed a set of strains each carrying a C-terminally ALFA-tagged POI. To rigorously test the live-cell imaging capabilities of ALIBY, we chose proteins with diverse spatio-temporal dynamics and different cellular functions. The tagging was confirmed by Western blot analysis using an HRP-conjugated anti-ALFA nanobody (sdAb) (**Fig. 2a**). Further, we performed immunoprecipitation (IP) for Gin4^ALFA^, which is a septin-associated polarity kinase(18), using the ALFA Selector^PE^ resin (**Fig. S1**) and were successful in detecting a sole specific band corresponding to Gin4^ALFA^ in the immunoprecipitated sample (expected molecular weight 130 kDa). This demonstrates that the tagging plasmids can be used to successfully tag yeast proteins at their C-terminal for biochemical applications.

**Fig. 2.**
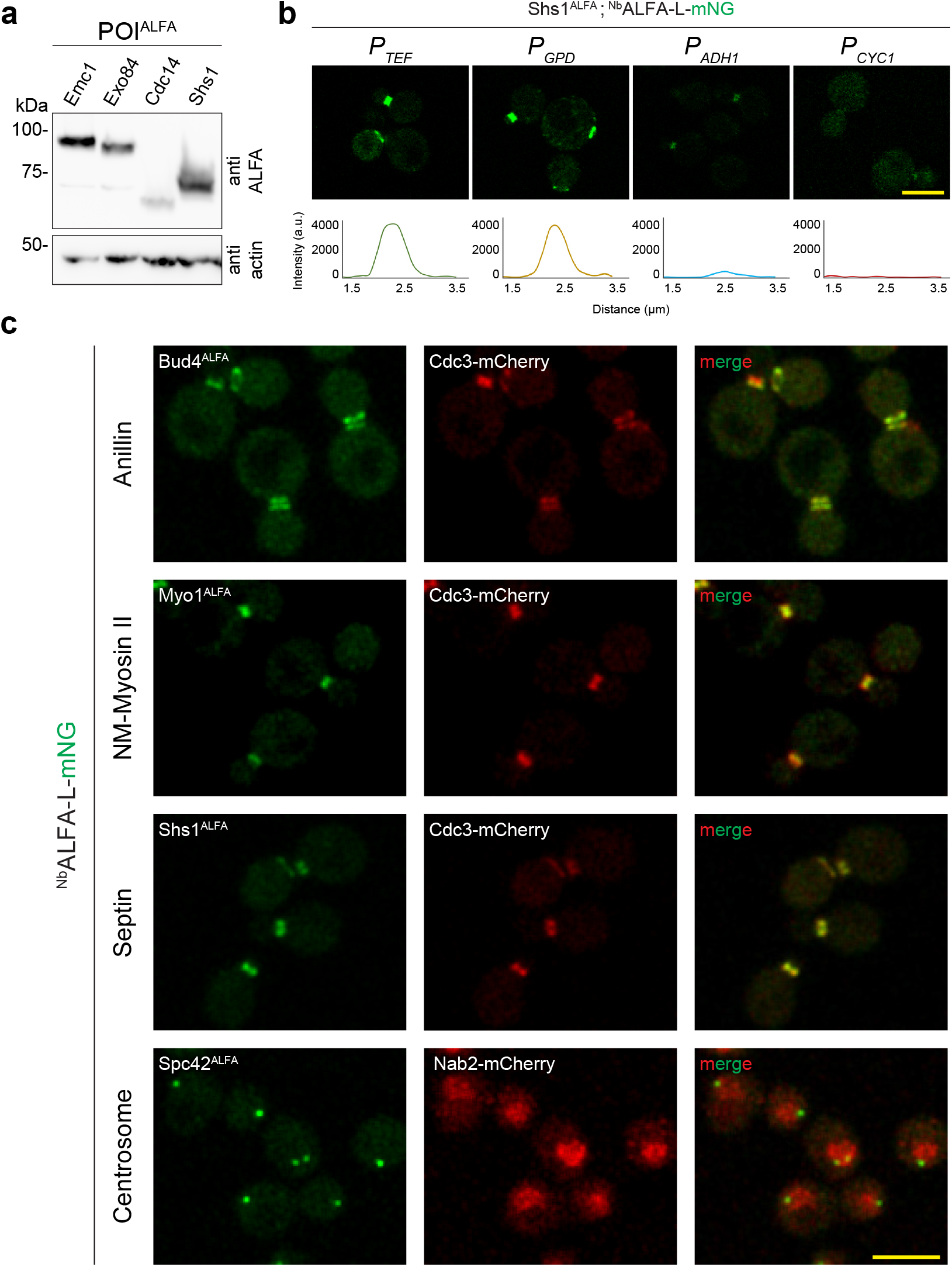
Characterization of plasmids and live-cell imaging of various proteins. **a** Confirmation of ALFA-tagged proteins by Western blotting. Whole cell extracts of the control strain lacking the ALFA tag and strains containing ALFA-tagged Emc1 (87 kDa), Exo84 (86 kDa), Cdc14 (62 kDa), and Shs1 (63 kDa) respectively, were treated with HU-DTT, resolved by SDS-PAGE, and analyzed by Western blot using anti-ALFA sdAb. The actin loading control was detected using a mouse actin mAb. **b** Comparison of promoters for optimizing ^Nb^ALFA expression by laser point scanning confocal microscopy: (Top row) ALFA-tagged Shs1 in ESM356 cells carrying *pRS305-promoter-* ^Nb^ALFA-L-CmNG-*term_CYC1_*; left to right: *P_TEF_, P_GPD_, P_ADH1_, P_CYCR_*. Images were captured at 1% laser excitation power at 488 nm. (Bottom row) Intensity profiles for cells with Shs1^ALFA^ expressing ^Nb^ALFA-L-mNG under different promoters in the same order as above. Scale bar: 5 μm. **c** Imaging of ALFA-tagged proteins: Left to right (in each row): ALFA-tagged POI along with ^Nb^ALFA-L-mNG under *P_TEF_* (green); mCherry-tagged marker protein (red); merged image. First row: Bud4, an anillin-like protein; Second row: Myo1, a myosin heavy chain; Third row: Shs1, a terminal septin. Cdc3-mCherry was used as a bud neck marker for the above ALFA-tagged proteins. Fourth row: Spc42, a spindle pole body component, with Nab2-mCherry as the nuclear marker. All the strains were grown in SC media at 30°C. Scale bar: 5 μm.

An ideal live-cell imaging setup demands a high signal-to-noise ratio at a low laser excitation to reduce phototoxicity while maintaining nanobody expression at a level such that normal functioning of the tagged protein is not hampered. Thus, to optimize ^Nb^ALFA expression for quantitative live-cell imaging, we integrated detection plasmids expressing the ^Nb^ALFA-L-mNG construct under different promoters into the strain carrying Shs1^ALFA^ (a terminal septin involved in cytokinesis(19)) and compared fluorescence signal strengths across the different promoters using laser point scanning confocal microscopy (**Fig. 2b**). Shs1 was chosen due to its exclusive localization to the bud neck which allows for easy quantitative analysis across the range of promoters. A qualitative comparison of relative signal strengths was first made at the same laser excitation power. We found that a strong fluorescence signal is achieved with nanobody expression under *P_GPD_* and *P_TEF_* promoters, accompanied with low cytoplasmic background, while in the case of *P_ADH1_* and *P_CYC1_* promoters, the fluorescence signal is weaker (**Fig. 2b**). For a quantitative comparison, we analyzed the intensity profiles for each promoter across a line scan parallel to the mother-bud axis. We found a large, clear peak in fluorescence intensity for *P_GPD_* and *P_TEF_*, while the intensity peaks for *P_ADH1_* and *P_CYC1_* are comparable to the baseline level (**Fig. 2b, S1b**). This set of observations implies that both *P_GPD_* and *P_TEF_* offer an excellent signal-to-noise ratio as compared to *P_ADH1_* and *P_CYC1_*. For *P_TEF_* and *P_GPD_*, the fluorescence signal was specific with minimal background even at low laser power, making them ideal candidates for use in live-cell imaging. Control strains expressing mNeonGreen without the ALFA tag or the ^Nb^ALFA expectedly showed a uniform cytoplasmic signal, lacking any foci corresponding to a localized protein (**Fig. S1c**). We therefore chose the *P_TEF_* promoter for nanobody expression in all subsequent experiments. To further verify the live-cell imaging capabilities of ALIBY, we imaged proteins localizing to various subcellular locations, such as Bud4 (anillin-like protein)(20), Myo1 (non-muscle type II myosin)(21), and Spc42 (component of spindle pole body (SPB)(22), the centrosome homolog in yeast), along with appropriate location markers (C-terminally mCherry-tagged Cdc3(23), a component of the septin complex, as a bud neck marker; C-terminally mCherry-tagged Nab2 as a nuclear marker(24)) (**Fig. 2c**). We obtained a similarly high signal-to-noise ratio without compromising protein localization, while also preserving normal cellular morphology, confirming the utility of ALIBY for live-cell imaging of various yeast proteins.

### Live-cell imaging of ALFA-tagged proteins

Next, to test the employability of ALIBY for live-cell imaging over extended durations, we proceeded to image ALFA-tagged proteins of varying abundances which localize to various subcellular localizations such as the bud neck, exocyst, SPB, and nucleus, using the ^Nb^ALFA-L-mNG construct expressed under the *P_TEF_* promoter.

To visualize the bud neck region and as a cell cycle marker, we used Cdc3-mCherry(23). We first imaged Bud4 (**Fig. 3a, Supplementary Video 1 and 2**), an anillin-like protein which marks the new bud-site(20), and associates closely with septins at the bud neck. It is also known to regulate the septin double ring structure(25). We then quantitatively compared the signal-to-noise ratios of Bud4^ALFA^ with conventional C-terminally GFP-tagged Bud4 (**Fig. 3d**). We also find that the spatiotemporal localization pattern of Bud4^ALFA^ exactly matches that of Bud4-GFP, as depicted by their kinetic profiles at the bud neck (**Fig. 3e**), implying that the cell cycle kinetics of Bud4 remain unperturbed upon ALFA-tagging as compared to a GFP tag. We then observed Bni5, which interacts with and provides stability to the septin ring at the mother-bud neck (**Fig. 3b, Supplementary Video 3 and 4**)(26). Bni5 is recruited to the bud neck at the early bud stage and leaves just before the septin hourglass-to-double ring (HDR) transition(27). The signal-to-noise ratio of Bni5^ALFA^ and its kinetics again closely match that of Bni5-GFP (**Fig. 3d, 3f**). Next, we visualized Shs1 (**Fig. 3c, Supplementary Video 5**), a terminal septin which is a member of the septin scaffold at the bud neck(19), which is essential for actomyosin ring (AMR) assembly and disassembly(28). Shs1 is also known to recruit Bni5 to the mother-bud neck(29). A similar analysis of Shs1^ALFA^ kinetics when compared to Cdc3-mCherry shows perfect synchronization between the two septins (**Fig. 3g**), which strengthens the finding that the localized accumulation kinetics of the above proteins during the cell cycle are preserved in the ALFA-tagged strains without any detectable off-targeting.

**Fig. 3.**
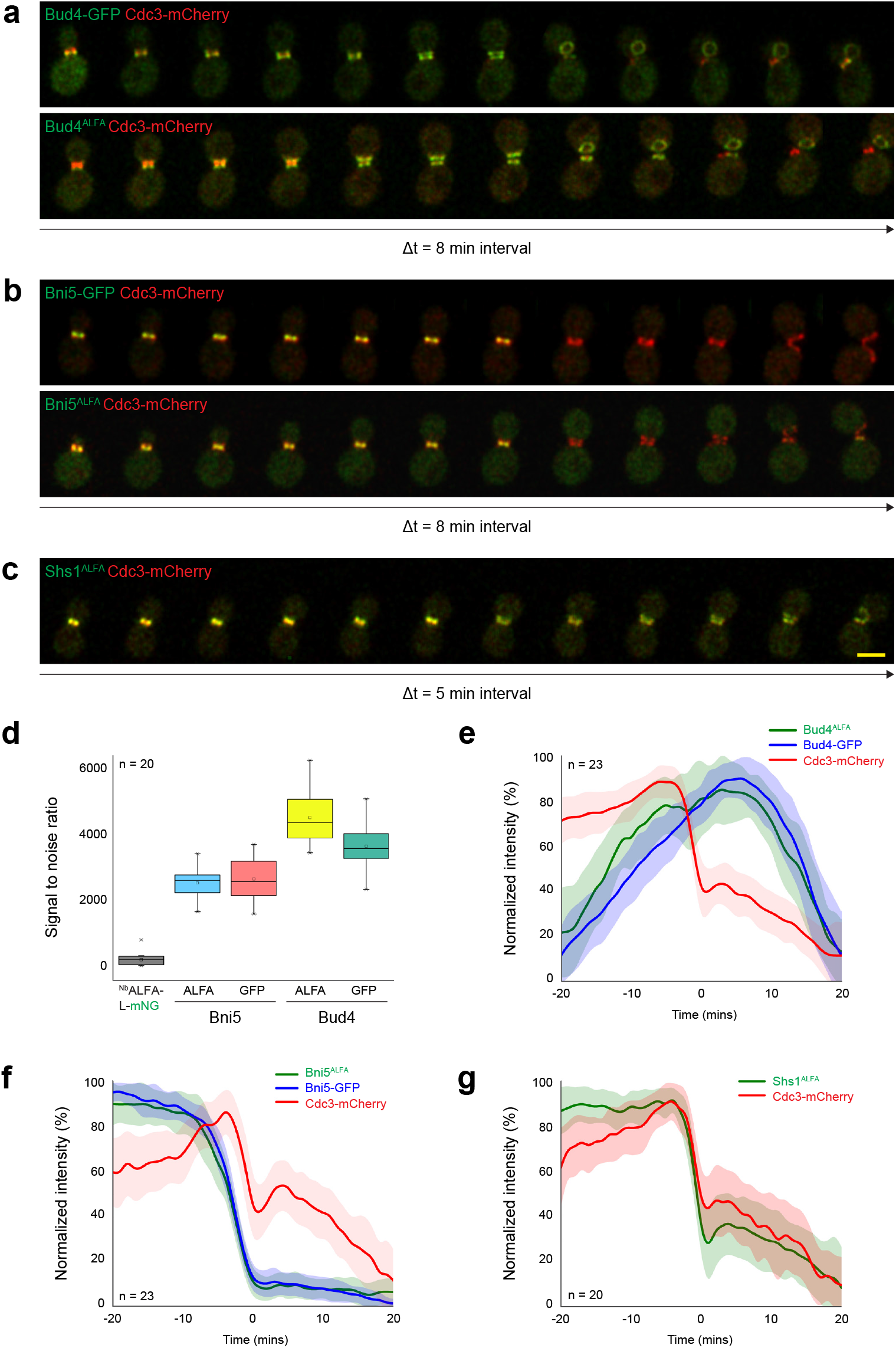
Timelapse imaging and quantification of protein kinetics. Time interval between successive images shown in each montage is denoted as Δt. **a** (Top) Bud4-GFP and (bottom) Bud4^ALFA^; Δt = 8 minutes. **b** (Top) Bni5-GFP and (bottom) Bni5^ALFA^; Δt = 8 minutes. **c** Shs1^ALFA^; Δt = 5 minutes. **d** Signal-to-noise ratio comparison between C-terminal ALFA tag and GFP fusions of Bud4 and Bni5 (n = 20). **e** Comparison between kinetics of Bud4-GFP and Bud4^ALFA^ (n = 23). **f** Comparison between kinetics of Bni5-GFP and Bni5^ALFA^ (n = 23). **g** Comparison between kinetics of Shs1-GFP and Cdc3-mCherry (n = 20). In addition to the ALFA-tagged proteins, the above ALFA tag containing strains also expressed ^Nb^ALFA-L-mNG under *P_TEF_.* All the strains were grown in SC media at 30°C. ALFA-tagged proteins bound to ^Nb^ALFA-L-mNG, and GFP-tagged proteins: green; Cdc3-mCherry: red. All the timelapse images were maximum intensity projected and deconvolved for representation in montages. Scale bar: 5 μm.

Next, we imaged proteins with diverse subcellular localizations, starting with Hof1, an F-BAR protein associated with the septin scaffold at the bud neck, which organizes actin filaments and is a part of the ingression-progression complex that coordinates primary septum formation with AMR constriction (**Fig. S2a**)(30). Myo1, a type II myosin heavy chain which contributes to the generation of force required for cytokinesis, is brought to the early bud-site by Bni5 (**Fig. S2b, Supplementary Video 6**)(26). Myo1 associates with F-actin throughout anaphase to give rise to the AMR, which constricts to separate the daughter cell from the mother cell(31). Next, we focused on visualizing proteins localizing to other subcellular locations. One such protein was Exo84, an important member of the exocyst complex involved in plasma membrane expansion and fusion through delivery of secretory vesicles to active sites of exocytosis. In nascent buds, Exo84 concentrates at the bud tip, following which it gradually diffuses, and during cytokinesis it shifts localization to the bud neck (**Fig. S2c, Supplementary Video 7**)(32). We then observed Cdc14, a phosphatase localized to the nucleolus (marked by Nab2-mCherry(14)) which gets released during anaphase through the MEN and FEAR pathways (**Fig. S2d, Supplementary Video 8**)(33, 34). Finally, we visualized Spc42, a constituent of the SPB. The SPB duplicates in G1/S phase and migrates to opposite ends of the mother nuclear envelope forming the mitotic spindle, and following cytokinesis, a copy of the SPB moves to the daughter cell (**Fig. S2e, Supplementary Video 9**)(22).

All the ALFA-tagged strains exhibited normal cell morphologies in our time-lapse studies and showed similar growth rates to the wild-type and the strains containing their GFP-tagged counterparts in a spot assay (**Fig. S2f**). We also find that the presence of the tag or the nanobody in isolation or in combination does not affect cellular growth across a wide range of temperatures. Furthermore, as discussed previously, we observe that the cell cycle kinetics across diverse proteins remain preserved upon fusion with a C-terminal ALFA tag. Thus, in summary, ALIBY offers an excellent signal-to-noise ratio for long-duration quantitative live-cell imaging of a variety of differentially abundant yeast proteins, while preserving normal cell physiology, growth rates, protein localization and kinetics.

## Discussion

The ALFA tag was designed and characterized as a competent, all-in-one tag with a unique sequence absent in commonly used eukaryotic model organisms which can be reliably used to study protein functions(3). This 13 amino acid tag possesses many desirable features such as versatility, compactness, high solubility, and the ability to refold upon stringent chemical treatments(3), which underscores the tag’s suitability for diverse applications such as live-cell imaging, immunoblotting, and immunoprecipitation, making the ALFA/^Nb^ALFA system a powerful tool for studying protein dynamics and function.

Here, we have extended the benefits of the ALFA/^Nb^ALFA system to yeast research through a toolkit comprising of a series of vectors, specifically designed for tagging and detecting any POI. These vectors are ready-to-use, requiring no further cloning steps, with several selectable markers to choose from, to introduce the ALFA tag or ^Nb^ALFA-L-mNG into the strain of interest. We have based our tagging plasmids on the well-established pYM series of vectors which rely on chromosomal integration of PCR amplified cassettes by homologous recombination(1). This makes the adaptability of ALIBY into the conventional yeast genetics workflow seamless and convenient, as the widely used long C-terminal tagging primers(13) (S2/S3) are compatible with our kit. Similarly, the detection plasmids build upon the commonly used pRS30-series of vectors and can be readily integrated into the genome after linearization with the same enzymes as used for the pRS series.

We have demonstrated the extensive set of applications of ALIBY through live-cell imaging and biochemical studies of various yeast proteins. Through our primary focus on live-cell imaging of numerous proteins localized at various subcellular locations, we have established the toolkit’s ability to derive an excellent signal-to-noise ratio with no detectable off-targeting for all proteins tested in our study at different subcellular locations, thus, making it suitable for quantitative imaging over an extended time duration, without affecting cellular growth and morphology. In addition, imaging of conventional fluorescent protein-tagged POIs over long periods of time inevitably brings with it the drawback of signal intensity loss due to photobleaching. ALIBY might not suffer from this drawback, since the high on- and off-rates of the ALFA/^Nb^ALFA complex(3) imply that ^Nb^ALFA-L-mNG molecules from the diffuse cytoplasmic pool can replace the POI^ALFA^-bound pool. This might effectively compensate for the photobleaching that takes place at the specific POI location during longer durations of live-cell imaging. ALIBY can also be employed for biochemical and cell biology applications such as coimmunoprecipitation (CoIP), mass spectrometry, immunofluorescence (IF), and super resolution microscopy, making it a valuable asset for the yeast research community. By simplifying the workflow for yeast protein investigations while employing the significant benefits of the ALFA tag over other epitope tags, we hope that ALIBY will contribute to accelerating yeast research and encourage innovative experimental designs to address a wider range of fundamental questions.

## Acknowledgements

We thank Prof. P. N. Rangarajan (Dept. of Biochemistry, IISc), Prof. Sunil Laxman (InStem, Bangalore), Prof. Gislene Pereira (CoS, University of Heidelberg) and Prof. Erfei Bi (University of Pennsylvania) for plasmids and reagents. This work was supported by SERB SRG grant (SRG/2021/001600), and Department of Biotechnology-Wellcome Trust India Alliance Intermediate fellowship (IA/I/21/1/505633) which was awarded to S.P. We acknowledge the DBT-IISc Partnership Program Phase-II (BT/PR27952/IN/22/212/2018), and the Department of Science and Technology, Ministry of Science and Technology, India DST-FIST Programme funded Confocal Microscopy Facility, Dept. of Biochemistry, IISc. D.A. acknowledges the KVPY (DST) fellowship program. A.D. acknowledges GATE fellowship from IISc. B.B. acknowledges the DST-INSPIRE fellowship program.

## Author contributions

S.P. conceptualized the overall project, provided resources and funding, and performed formal analysis. D.A., A.D., and A.S.V. conducted the experiments and collected data. B.B. performed image analysis. D.A., A.D., A.S.V., and S.P. wrote the manuscript and contributed to the editing and review of the final manuscript.

## Materials and methods

### Cloning and construction of plasmids

All plasmids used in this study are listed in Supplementary Table 1, and the primers used for cloning the constructs and tagging proteins-of-interest are listed in Supplementary Table 2. For constructing the tagging plasmids, oligos carrying the ALFA tag sequence codon optimized for yeast and flanked by SalI and BglII sites were synthesized by Sigma-Aldrich. These oligos were annealed in annealing buffer (10 mM Tris pH 7.5-8.0, 50 mM NaCl, 1 mM EDTA) as recommended by Addgene (https://www.addgene.org/protocols/annealed-oligo-cloning), then digested with SalI and BglII, followed by ligation with SalI-BglII digested pYM14/16/17 and *pFA6a-HIS3MX6* to give rise to tagging plasmids piSP532/534/536/538.

For constructing the detection plasmids, a sequential strategy was adopted. Promoter-MCS-*term_CYC1_* fragments were PCR amplified from pRS414/415 vectors carrying *P_CYC1_/P_ADH1_/P_GPD_/P_TEF_* promoters. These fragments were cloned into pRS305 (linearized with XbaI and XhoI) using NEBuilder^®^ HiFi DNA Assembly as per the manufacturer’s instructions (Cat. No. E2621, New England BioLabs). These were subsequently transformed into *E. coli* TOP10 cells, plasmids were isolated using QIAprep Spin Miniprep Kit (Cat. No. 27106, QIAGEN) and confirmed by restriction digestion. The ^Nb^ALFA-L-mNG sequence was synthesized and cloned as mentioned into the above vectors (linearized with BamHI) to give rise to piSP551/553/555/557 respectively, which are our detection plasmids. Based on imaging results, the ^Nb^ALFA-L-mNG construct alongwith *P_TEF_* and *term_CYC1_* was then cloned as previously into pRS303/304/306 (linearized with BamHI) to generate piSP765/piSP767/piSP565 respectively, which are the rest of our detection plasmids. For constructing the control plasmid piSP763, the L-mNG fragment was amplified from piSP557 and cloned into BamHI digested piSP525. piSP629 was constructed by cloning mCherry (amplified from mCherry-pBAD, Addgene #54630) into pYM17 (digested with SalI and BglII). All the above plasmids were confirmed by insert release using restriction digestion and sequencing, and will be submitted to Addgene as the ALIBY toolkit.

### Construction of yeast strains

All yeast strains constructed for this study are listed in Supplementary Table 3. The wild-type strain used was *Saccharomyces cerevisiae* ESM356, derived from the S288C genetic background. Yeast strains were cultured at 30°C and 180 rpm rotation(35). For yeast strain construction, we used YPD agar plates supplemented with antibiotics – 100 μg/mL G418 (Cat. No. 58327, Sisco Research Laboratories Pvt. Ltd.), 100 μg/mL nourseothricin (Cat. No. AB-102, Jena Bioscience), 50 μg/mL hygromycinB (Cat. No. 67317, Sisco Research Laboratories Pvt. Ltd.) or synthetic complete (SC) medium agar plates lacking Leucine or Tryptophan, and corresponding liquid media.

In order to introduce the bud neck marker, plasmid pMO014 was linearized with BglII (Cat. No. R0144, New England BioLabs) and transformed for integration into the genomic *trp1* locus using a LiOAc-based protocol(14). To introduce the nucleus marker, PCR amplification was done from plasmid piSP629 using S2/S3 primers for Nab2, and the linear fragment was then transformed to obtain a C-terminal mCherry fusion of Nab2. For fusing the ALFA tag to the C-terminal of a desired set of proteins, PCR amplification was done from piSP534 using appropriate S2/S3 primers, and the linear fragments were transformed into appropriate strains containing corresponding location markers. For integration of the C-terminally mNeonGreen-fused ^Nb^ALFA into the genome, piSP522/523/524/525 were linearized with KasI (Cat. No. R0544, New England BioLabs) and transformed into the above strains as appropriate. To construct strain YSP210, plasmid piSP763 was linearized with KasI and transformed into the wild-type ESM356 strain YSP002. The PCR fragment for ALFA-tagging Shs1 generated using S2/S3 primers was then transformed into YSP210 to give rise to strain YSP248. To construct strains YSP126 and YSP286, pMO014 was linearized with BglII and transformed into YSP002. PCR amplified fragments from pYM25 using S2/S3 primers for Bud4 and Bni5 respectively were transformed into the above strain to obtain C-terminal GFP fusions.

### Spot assay

Yeast strains were pre-cultured in YPD broth overnight at 30°C, 180 rpm rotation, and subcultured to a final OD of 1. Serial dilutions 10^0^, 10^-1^, 10^-2^, 10^-3^, and 10^-4^ were made in YPD and 4 μL of each dilution was spotted onto 3 YPD agar plates. The plates were incubated at 23°C, 30°C and 37°C, respectively, for 48 hrs after which the plates were scanned.

### Live-cell imaging

For tethering yeast cells to glass-bottomed dishes (35mm, Cat. No. 81218-200, ibidi GmbH) for live-cell imaging, we coated them with 6% concanavalin A type 6 (Cat. No. C2010, Sigma-Aldrich). Yeast cells were pre-cultured overnight and were sub-cultured for 3-4 hrs to mid-log phase (OD 0.5-0.8) in SC broth at 30°C and 180 rpm rotation and plated on concanavalin A coated dishes. Live-cell imaging was performed using laser point scanning confocal microscope Olympus FV3000 equipped with 100x oil objective, high sensitivity GaAsP PMT detectors, solid-state lasers (488 nm and 561 nm). Images were acquired using Olympus FluoView 3000 (2.4.1.198) software. All timelapse movies consisted of 7 z-slices at a step size of 0.5μm, for a period of 60-180 mins with a 1-minute interval. 3D Deconvolution of raw images was done using Olympus CellSens Dimension (3.1) software followed by a maximum intensity projection using Fiji for representation in a montage. Signal-to-noise ratios were calculated on sum intensity projected raw images using Fiji as described previously(36). Quantification of protein kinetics was also done on sum intensity projected raw images using Fiji as described previously(37). Data was plotted in OriginPro 9.0 (OriginLab).

### Western blotting

Yeast cultures were grown in YPD broth till mid-log phase at 30°C and cell lysates were prepared using a trichloroacetic acid (TCA) based protocol as described(13). To summarize, yeast cells corresponding to 2 mL cultures were pelleted at 20000 *g* (full speed) for 3 mins and after removal of the supernatant, were resuspended in 800 μL cold water and vortexed. Next, they were incubated on ice with 150 μL of 1.85 M NaOH for 5 mins, followed by incubation on ice with 150 μL 55% TCA for 20 mins. The lysates were then pelleted at full speed for 20 mins and the supernatant was removed. The pellet was resuspended in 75 μL HU-DTT (8 M urea, 5% w/v SDS, 200 mM Tris-Cl pH 6.8, 0.1 mM EDTA, bromophenol blue, 15 mg/mL DTT) and vortexed, followed by neutralization with 1 μL 2 M Tris. The samples were then incubated at 95°C for 5 mins and centrifuged at full speed for 2 mins. 10-30 μL of the supernatants (based on protein abundances) were run on an 8% SDS-PAGE gel and transferred to a nitro-cellulose membrane (Cat. No. 1620112, BioRad) using a semi-dry transfer method (Bio-Rad, at 0.11 A limited to 16 V for 1.5 hrs) for Western blotting. The membrane was blocked with 3% skimmed milk in 1X PBS-T for 1 hr at room temperature followed by incubation with a a 1:1000 dilution of HRP-conjugated sdAb anti-ALFA primary antibody (Cat. No. N1505, Lot No. 15201101, NanoTag Biotechnologies) overnight at 4°C.

The membrane was then washed thrice with 1X PBS-T, and the blot was developed using ECL substrate (Cat. No. 170-5060, Bio-Rad). Images were acquired using ImageQuant 500 (GE). The same blot was stripped with stripping buffer (0.2 M glycine, 0.1% w/v SDS, 1% v/v Tween 20, pH adjusted to 2.2) for 30 mins and blocked again. Western blotting was performed to detect the actin loading control using a 1:500 dilution of mouse anti-actin monoclonal primary antibody (Cat. No. MA1-744, Lot No. WC317616, Invitrogen) and 1:5000 dilution of HRP-conjugated rabbit anti-mouse secondary antibody (Cat. No. 7076, Lot No. 36, Cell Signaling Technology), and the blot was developed similarly.

### Immunoprecipitation

Yeast cultures were grown in YPD broth till log phase (OD = 1) at 30°C and total cell extracts were prepared as described previously(38), modified for our purpose. To summarize, yeast cells corresponding to 50 mL cultures were pelleted at 2000 *g* for 5 mins (all centrifugation steps were performed at 4°C), and the pellets were washed with 1X PBS. After removal of the supernatant, the pellet was resuspended in 250 μL of lysis buffer (50 mM Tris-HCl pH 7.5, 120 mM KCl, 5 mM EDTA, 0.1% Nonidet P-40, 10% glycerol, 1 μL/mL DTT, 10 μL/mL PMSF, 1X PIC) and transferred to a 2 mL conical FastPrep vial to which an equal volume of chilled, acid-washed glass beads was added. Cell lysis was carried out at 4°C using a FastPrep-24™ Cell Disruptor (Cat. No. 6004500, MP Biomedicals) set at 6.5 for 8 cycles of 1 min each, with 2 mins of incubation in an ice water bath between cycles. Lysis was verified using Trypan Blue staining. Caps of the FastPrep vials were perforated using a sterile needle, and they were inverted into 15 mL conical tubes. The lysate was collected by centrifugation at 1000 *g* for 1.5 mins and transferred to a 1.5 mL microcentrifuge tube. Lysates were clarified by centrifuging at 16000 *g* for 10 mins and collecting 220 μL of the supernatant, and this step was repeated twice. 20 μL of the supernatant (1:10) was aliquoted as total cell extract (TCE) samples, and protein concentration was estimated using a standard Bradford Assay. The rest of the clarified lysates (corresponding to 1 mg of total protein) was incubated with 20 μL of washed ALFA Selector^PE^ bead slurry (Cat. No. N1510, NanoTag Biotechnologies) at 4°C for 2 hrs with head-over-tail rotation. The beads were sedimented by centrifugation at 1000 *g* for 1.5 mins, and the supernatant was removed. The beads were then washed thrice with 1X PBS + 0.1% NP-40. The TCE and bead samples were incubated with 5X SDS sample loading buffer (5% w/v SDS, 25% v/v glycerol, 1 M Tris-Cl pH 6.8, 0.05% w/v bromophenol blue, 1% v/v β-mercaptoethanol) at 95°C for 10 mins, and then centrifuged at full speed for 2 mins. 4 μL (1:50) and 20 μL of the supernatants of TCE and bead samples respectively were run on an 8% SDS-PAGE gel. Western Blotting was performed using the protocol mentioned above.

---

**Supplementary Table 1:**
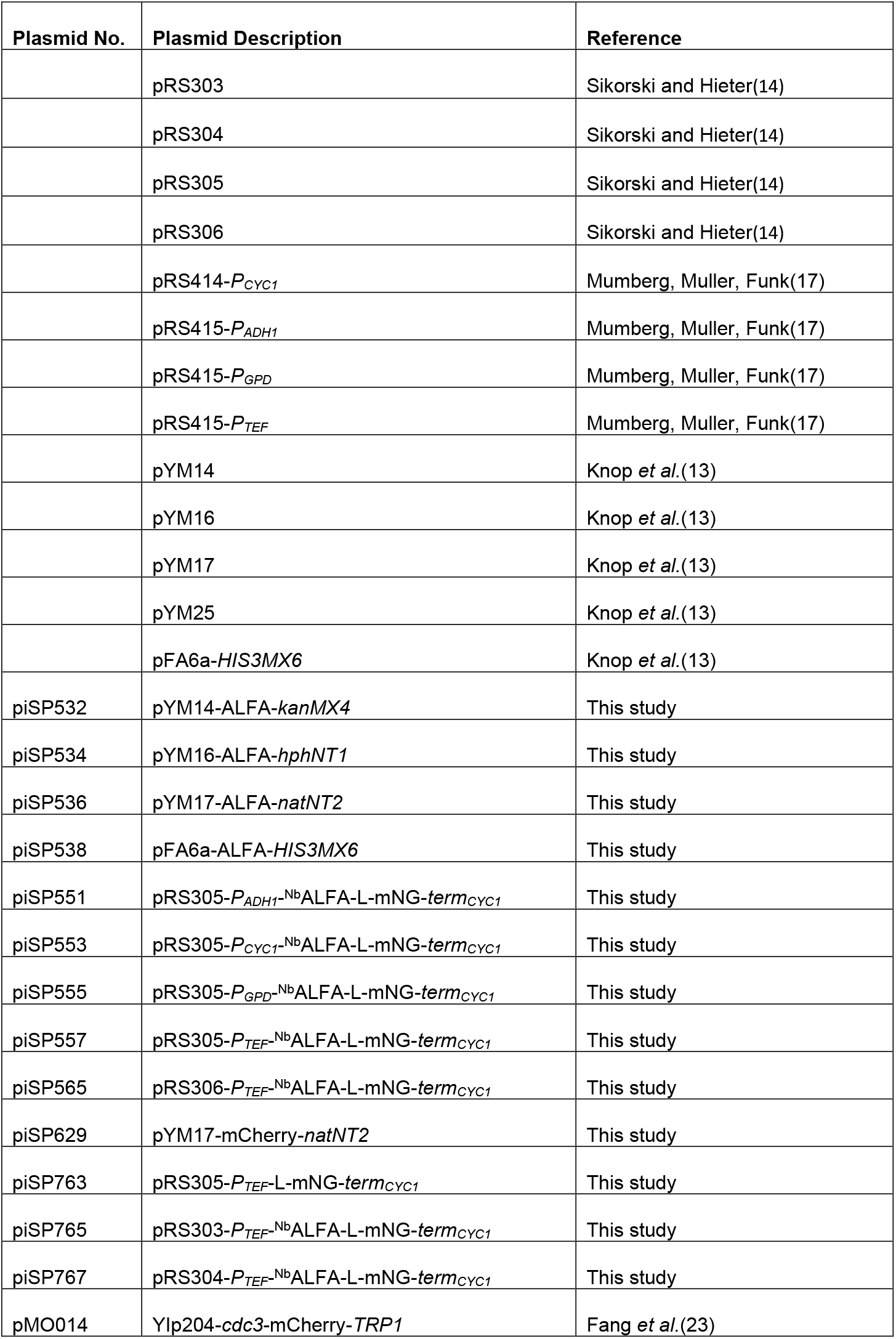
Plasmids used in this study.

**Supplementary Table 2:**
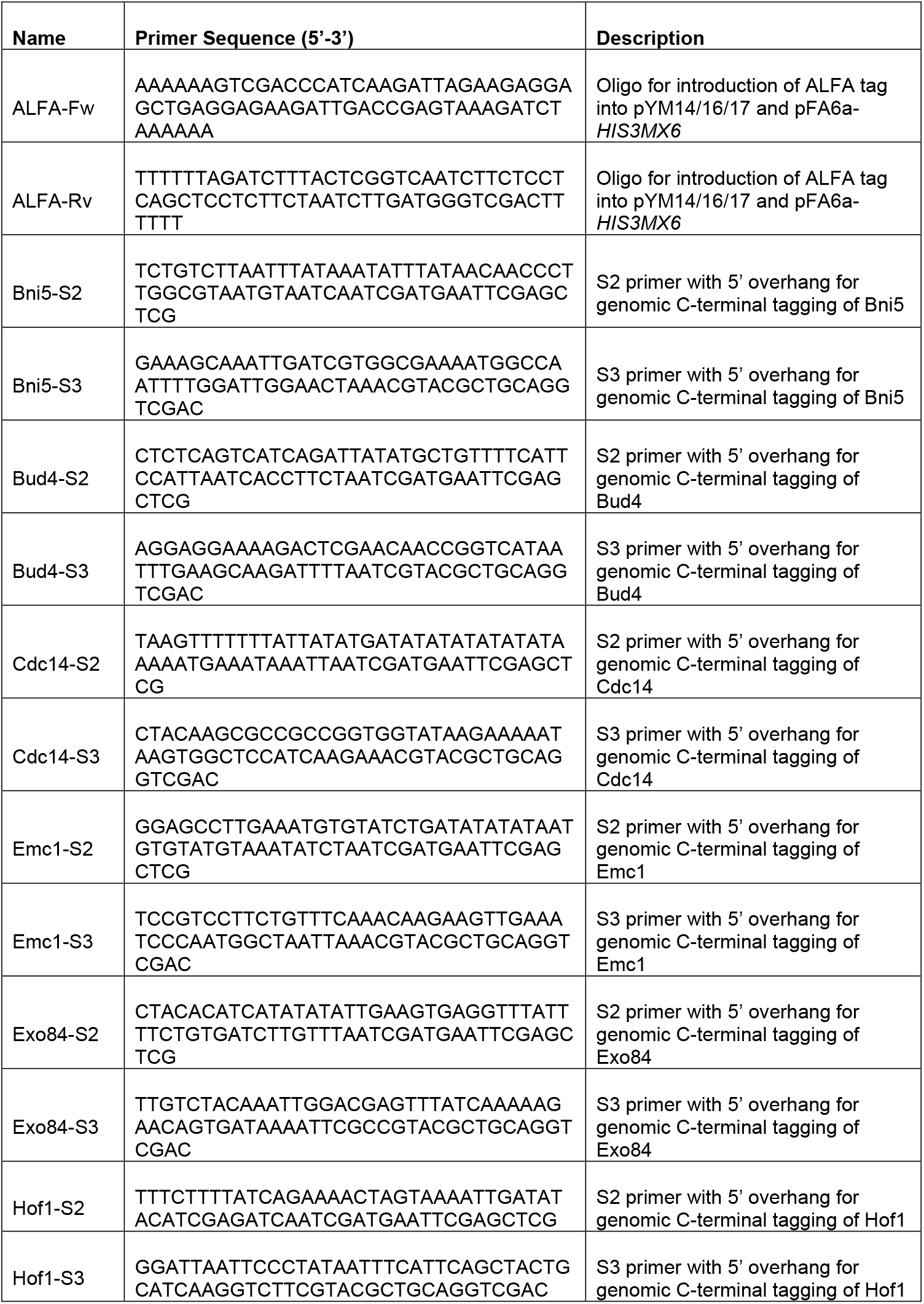
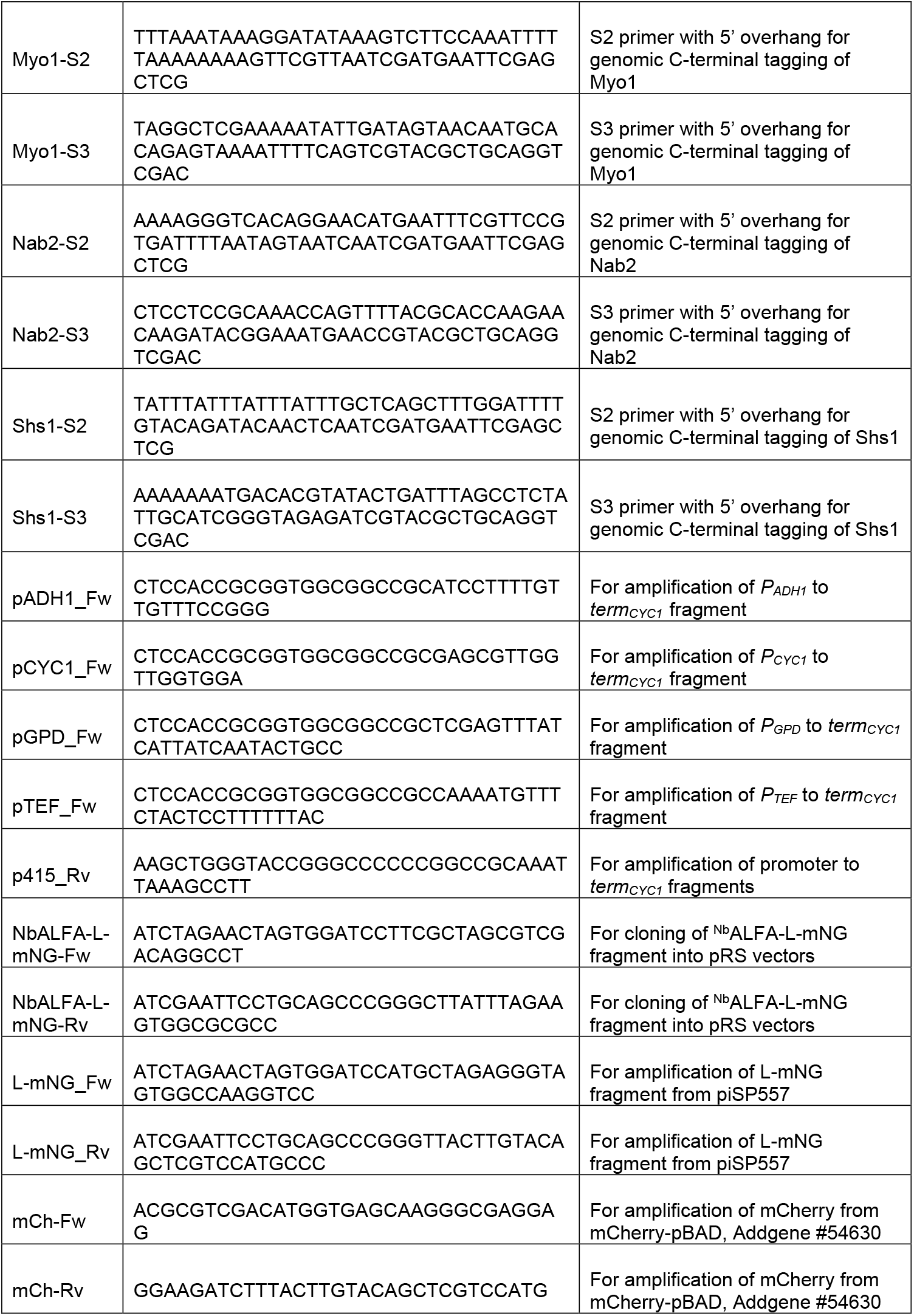
Primers used in this study.

**Supplementary Table 3:** Yeast strains used in this study.

| Strain No. | Description |
| --- | --- |
| YSP002 | ESM356 (*MATa ura3-52 leu2Δ1 trp1Δ63 his3Δ200*) WT |
| YSP126 | YSP002 *trp1<sup>+</sup>::cdc3-mCherry bud4-GFP::hphNT1* |
| YSP128 | YSP002 *leu2<sup>+</sup>::pRS305-P<sub>ADH1</sub><sup>Nb</sup>ALFA-L-mNG-term<sub>CYC1</sub>* |
| YSP130 | YSP002 *leu2<sup>+</sup>::pRS305-P<sub>CYC1</sub><sup>Nb</sup>ALFA-L-mNG-term<sub>CYC1</sub>* |
| YSP132 | YSP002 *leu2<sup>+</sup>::pRS305-P<sub>GPD</sub><sup>Nb</sup>ALFA-L-mNG-term<sub>CYC1</sub>* |
| YSP134 | YSP002 *leu2<sup>+</sup>::pRS305-P<sub>TEF</sub><sup>Nb</sup>ALFA-L-mNG-term<sub>CYC1</sub>* |
| YSP135 | YSP134 *trp1<sup>+</sup>::cdc3-mCherry bud4-ALFA::hphNT1* |
| YSP136 | YSP134 *trp1<sup>+</sup>::cdc3-mCherry myo1-ALFA::hphNT1* |
| YSP137 | YSP134 *trp1<sup>+</sup>::cdc3-mCherry exo84-ALFA::hphNT1* |
| YSP138 | YSP134 *trp1<sup>+</sup>::cdc3-mCherry* |
| YSP167 | YSP134 *nab2-mCherry::natNT2* |
| YSP169 | YSP167 *cdc14-ALFA::hphNT1* |
| YSP170 | YSP167 *spc42-ALFA::hphNT1* |
| YSP171 | YSP134 *trp1<sup>+</sup>::cdc3-mCherry shs1-ALFA::hphNT1* |
| YSP172 | YSP134 *trp1<sup>+</sup>::cdc3-mCherry bni5-ALFA::hphNT1* |
| YSP173 | YSP134 *emc1-ALFA::hphNT1* |
| YSP201 | YSP134 *trp1<sup>+</sup>::cdc3-mCherry hof1-ALFA::hphNT1* |
| YSP210 | YSP002 *leu2<sup>+</sup>::pRS305-P<sub>TEF</sub>-L-mNG-term<sub>CYC1</sub> trp1<sup>+</sup>::cdc3-mCherry* |
| YSP248 | YSP210 *shs1-ALFA::hphNT1* |
| YSP286 | YSP002 *trp1<sup>+</sup>::cdc3-mCherry bni5-GFP::hphNT1* |
| YSP299 | YSP002 *gin4-ALFA::hphNT1* |

**Supplementary Video Legends:**

**Supplementary Video 1.** (Left) Bud4^ALFA^; (middle) Cdc3-mCherry; (right) merge; Δt = 1 minute. Scale bar: 5 μm.

**Supplementary Video 2.** (Left) Bud4-GFP; (middle) Cdc3-mCherry; (right) merge; Δt = 1 minute. Scale bar: 5 μm.

**Supplementary Video 3.** (Left) Bni5^ALFA^; (middle) Cdc3-mCherry; (right) merge; Δt = 1 minute. Scale bar: 5 μm.

**Supplementary Video 4.** (Left) Bni5-GFP; (middle) Cdc3-mCherry; (right) merge; Δt = 1 minute. Scale bar: 5 μm.

**Supplementary Video 5.** (Left) Shs1^ALFA^; (middle) Cdc3-mCherry; (right) merge; Δt = 1 minute. Scale bar: 5 μm.

**Supplementary Video 6.** (Left) Myo1^ALFA^; (middle) Cdc3-mCherry; (right) merge; Δt = 1 minute. Scale bar: 5 μm.

**Supplementary Video 7.** (Left) Exo84^ALFA^; (middle) Cdc3-mCherry; (right) merge; Δt = 1 minute. Scale bar: 5 μm.

**Supplementary Video 8.** (Left) Cdc14^ALFA^; (middle) Nab2-mCherry; (right) merge; Δt = 1 minute. Scale bar: 5 μm.

**Supplementary Video 9.** (Left) Spc42^ALFA^; (middle) Nab2-mCherry; (right) merge; Δt = 1 minute. Scale bar: 5 μm.

**Fig. S1.**
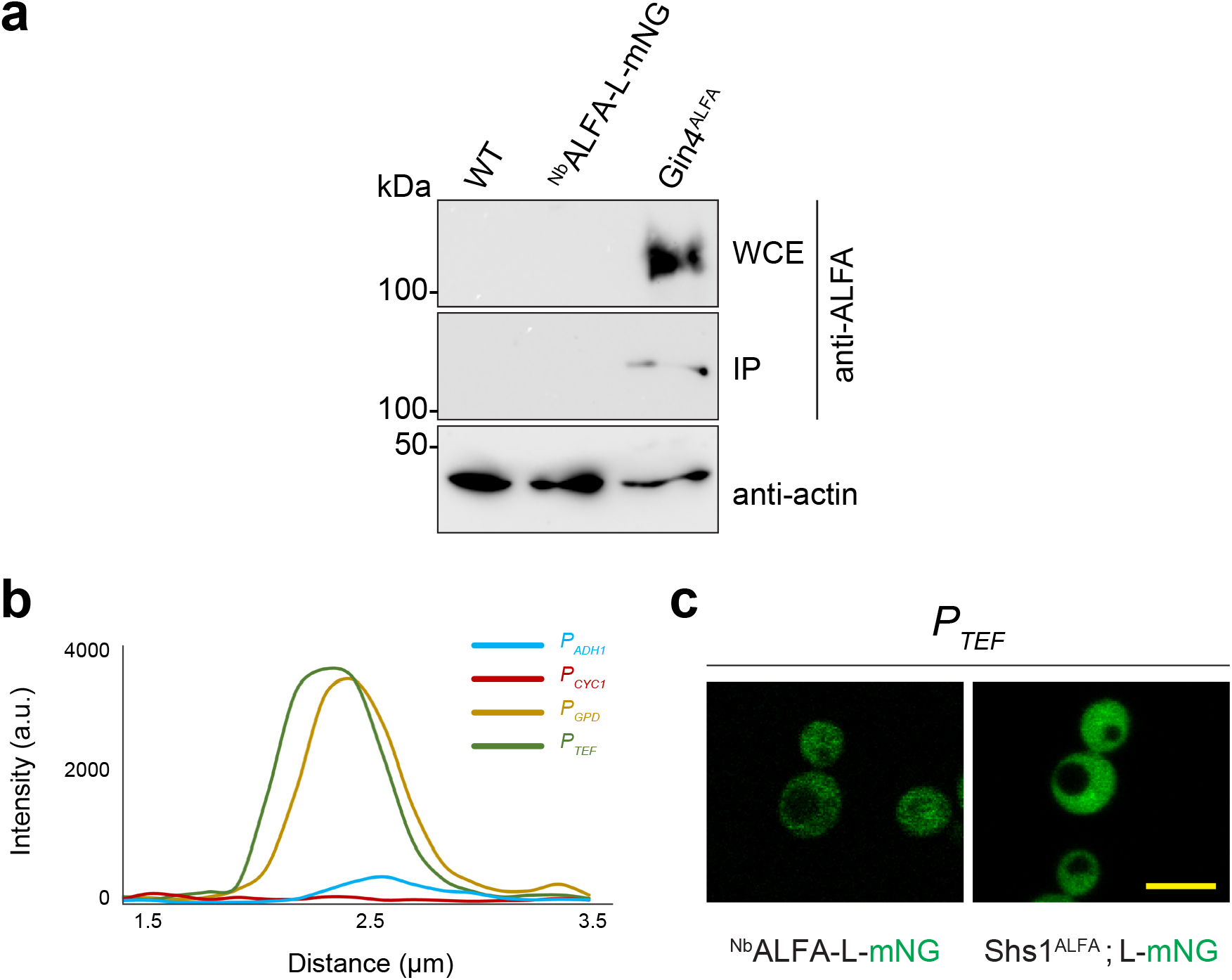
Characterization of plasmids. **a** Immunoprecipitation of Gin4^ALFA^ (expected molecular weight 130 kDa); the wild-type strain and the nanobody control strain lack the ALFA tag. Whole cell extracts (WCE) of each strain were incubated with ALFA Selector^PE^ resin and washed; 1/50 of the input fraction and the entire eluate fraction were resolved by SDS-PAGE and analyzed by Western blot using anti-ALFA sdAb. **b** Comparison between intensity profiles of line scans performed for the four promoters. **c** Laser point scanning confocal microscopy for further characterization. Control strains: (i) ^Nb^ALFA-L-mNG expressed without presence of an ALFA tag (ii) L-mNG without ^Nb^ALFA expressed in presence of Shs1^ALFA^

**Fig. S2.**
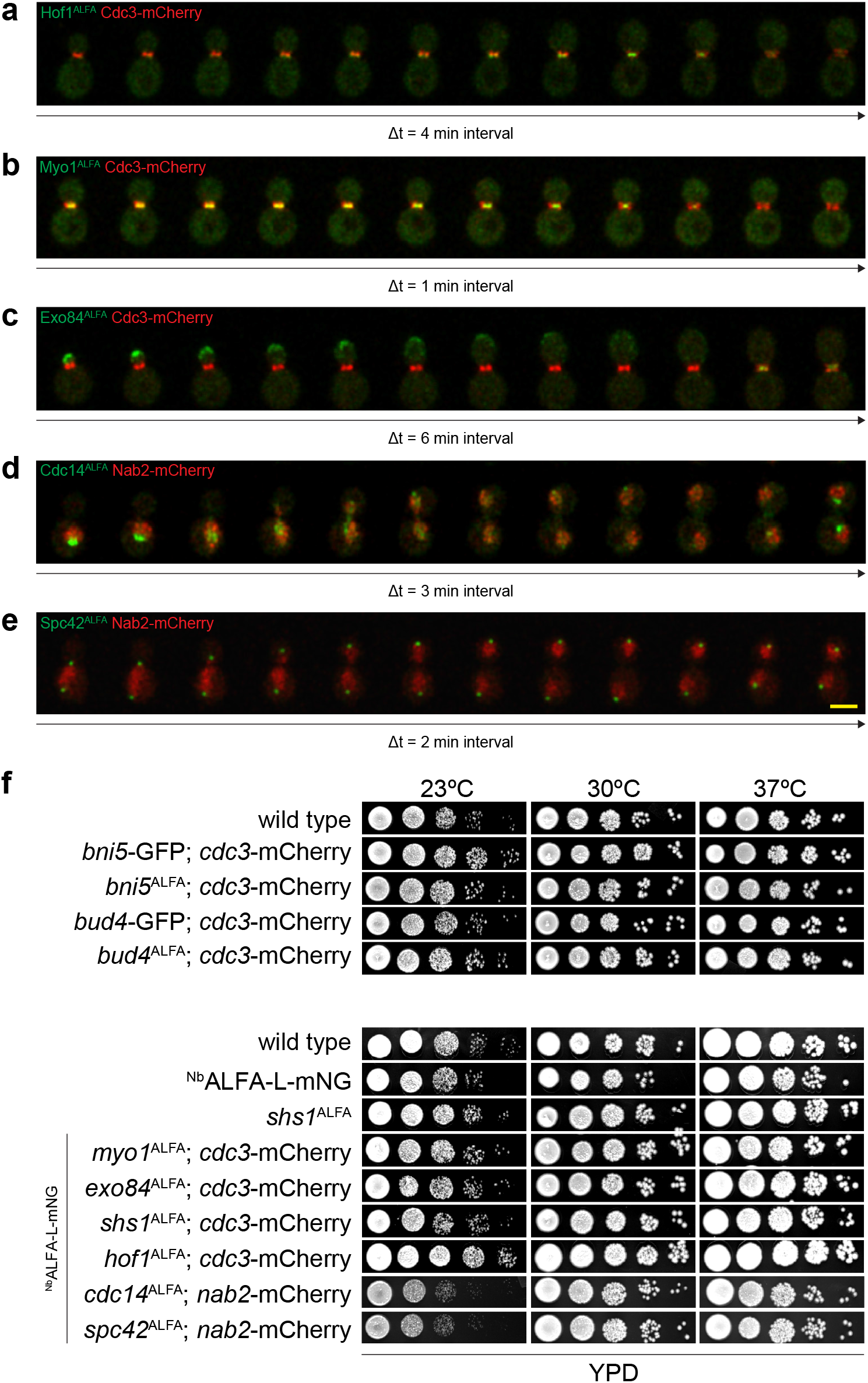
Timelapse imaging and spot assays. Time interval between successive images shown in each montage is denoted as Δt. **a** Hof1^ALFA^; Δt = 4 minutes. **b** Myo1^ALFA^; Δt = 1 minute. **c** Exo84^ALFA^; Δt = 6 minutes. **d** Cdc14^ALFA^; Δt = 3 minutes. **e** Spc42^ALFA^; Δt = 2 minutes. In addition to the ALFA-tagged proteins, the above strains also expressed ^Nb^ALFA-L-mNG under *P_ŢEF_.* All the strains were grown in SC media at 30°C. ALFA-tagged proteins bound to ^Nb^ALFA-L-mNG: green; mCherry-tagged marker proteins (Cdc3 for panels **a-c**, Nab2 for panels **d-e**: red. All the timelapse images were maximum intensity projected and deconvolved for representation in montages. Scale bar: 5 μm. **f** Comparison of growth rates of strains containing POI^ALFA^ with wild type strain, control strains and POI-GFP containing strains. Five serial dilutions for each strain were spotted onto YPD plates which were grown at 23°C, 30°C and 37°C.

